# Host plant use is driven by microclimate not nutritional quality in a grassland butterfly

**DOI:** 10.1101/2025.11.15.688506

**Authors:** William B.V. Langdon, Richard Fox, Owen T. Lewis

## Abstract

1. Abundance of insect herbivores often depends on host plant suitability for their specialised immature stages. Suitability can be strongly influenced by both microclimate and the nutritional quality of the plants themselves. Where soil nitrogen is high, host plants tend to have high nutritional quality, but vigorous growth of surrounding vegetation reduces microclimatic temperatures. Thus, thermophilous insects may face a choice between host plants with optimal microclimates and those with optimal nutritional quality.
2. We investigated how microclimate and nitrogen content influence oviposition choices by the declining Small Copper butterfly, *Lycaena phlaeas,* on its host plant, *Rumex acetosa*. We predicted that warmer plants would have lower nitrogen content, and that butterflies would choose cooler, high-nitrogen plants during warmer ambient conditions.
3. Although warmer *R. acetosa* plants had lower nitrogen content, *L. phlaeas* consistently chose to lay eggs on plants in warm microclimates, implying a trade-off between temperature and the nutritional quality of host plants.
4. Patches of bare ground created by *Talpa europaea* (European Mole) near *R. acetosa* plants increased microclimatic temperatures and decoupled the negative correlation between nutritional quality and thermal suitability.
5. Our results have implications for the conservation of thermophilous insect herbivores, especially close to their range margins and in the context of climate change. Rather than maximising host plant abundance or nutritional quality, management that creates suitable microclimatic conditions is likely to be critical. Our findings also suggest that, while nitrogen pollution may increase host plant nutritional quality, its negative impacts on microclimate will likely further reduce breeding habitat for *L. phlaeas* and other insects in grassland habitats.

## INTRODUCTION

The quality of habitat for insects is primarily dictated by the availability of sites in which the least mobile and most specialised early stages can survive (Bourn & Thomas, 2002; Thomas et al., 2011). For many herbivorous insects of open habitats, a key factor is the availability of suitably warm microhabitats around host plants, particularly for species close to their cool range margins (Oliver et al., 2009; Thomas et al., 1999). Here, ambient temperatures are often lower than those required for these insects to complete their development (van Wingerden et al., 1991). With limited ability to thermoregulate behaviourally (Ashe-Jepson et al., 2023) or move between feeding and thermoregulating sites, this means that immature stages can often only survive on host plants growing in particularly warm microhabitats (Curtis & Isaac, 2015; WallisDeVries, 2006). As a result, the availability of host plants in such microhabitats often dictates the abundance of thermophilous insect species (Curtis & Isaac, 2015; J.A. Thomas et al., 2001).

Microhabitat aside, not all host plants are of equal nutritional quality for insects, because of high inter- and intra-specific variability in plant chemistry (Persson et al., 2010). Insects require a wide variety of nutrients, each in specific amounts (Raubenheimer et al., 2009; Raubenheimer & Simpson, 1993, 2004). When the chemistry of food does not match these requirements, insects are forced to overeat and excrete excess nutrients to make up for deficiencies of others, at a cost of time and energy (Raubenheimer et al., 2005; Simpson et al., 2004). Thus, host plant nutritional quality is defined as the match between insect requirements and host chemistry (Sterner & Elser, 2002; Vogels et al., 2023), and large mis-matches can have strong effects on fitness that scale up to population-level impacts (Cease et al., 2012; Clissold et al., 2006).

Insects are, therefore, strongly constrained by their thermal and nutritional requirements, and their fitness will be highest on host plants where both of these are optimal. Such conditions may be rare, however, if thermally and nutritionally suitable conditions tend not to overlap. For example, the host plants of many herbivorous insects have a much higher carbon content and a much lower nitrogen content than the insects themselves (Elser et al., 2000). Thus, to attain sufficient nitrogen to complete their development, insects must generally overeat large quantities of carbon. Plant nitrogen content is, therefore, often regarded as a key indicator of host nutritional quality for insects, with performance generally improved on higher-nitrogen host plants (Mattson, 1980; Throop & Lerdau, 2004; White, 1993). A wide variety of factors can influence host plant nitrogen content, including the biology of the plant species concerned (for example an ability to fix nitrogen), its developmental stage (Thurner et al., 2025; Zhang et al,, 2013), or environmental conditions such as temperature, water or phosphorus availability, via their impacts on nitrogen uptake (Bista et al., 2013; Joseph et al., 2021; Ercoli et al., 1996). Nevertheless, in general, plant nitrogen content usually reflects soil nitrogen levels (Fujita et al., 2013; Sardans et al., 2012; Throop & Lerdau, 2004). In open habitats, however, higher soil nitrogen also enhances vegetation productivity, leading to taller vegetation, with reduced penetration of sunlight to ground level (Bobbink, 1991; DeMalach et al., 2017; Hautier et al., 2009). Microhabitat temperatures in such situations are lower than in more open locations where bare ground and dead plant material exposed to sunlight warm up rapidly and radiate heat (Curtis & Isaac, 2015; WallisDeVries, 2006). Therefore, in general, host plants with higher nutritional quality (nitrogen content) tend not to overlap with warm microhabitats, creating a potential trade-off for thermophilous insects in open habitats.

Such trade-offs have important implications for the effective conservation of insect populations, especially in the context of long-term declines in abundance of many terrestrial insect taxa around the world (Edwards et al., 2025; Van Klink et al., 2020; Warren et al., 2021). Both local habitat management and wider-scale policy instruments can influence insects’ exposure to changes in microclimate and nitrogen availability, so understanding species’ responses to these environmental variables will directly inform effective conservation. Furthermore, anthropogenic climate change and increasing diffuse nitrogen pollution are acknowledged global and regional drivers of insect population change (Carvalheiro et al., 2020; Hill et al., 2021; Roth et al., 2021; Wilson & Fox, 2021).

The roles of thermal and nutritional requirements in dictating insect microhabitat use are rarely studied simultaneously, however (e.g. Bourn & Thomas, 1993; Ellis, 2003). As a result, it is difficult to understand the relative strength of thermal and nutritional constraints for many insect herbivores, and whether our hypothesised trade-off exists. Such an understanding is important when climate change and nitrogen pollution are altering the thermal and nutritional conditions experienced by insects simultaneously (Rashid et al., 2023). Rising ambient temperatures under climate change may lessen thermal constraints on some insect species, as microhabitats that were previously too cool become sufficiently warm (Roy & Thomas, 2003; C.D. Thomas et al., 2001). On the other hand, warming extends the growing season (Menzel et al., 2020), enhancing plant productivity and potentially leading to cooler microhabitats overall (WallisDeVries & van Swaay, 2006). Rising levels of soil nitrogen under nitrogen pollution are expected to enhance plant productivity and alter vegetation structure alongside warming (WallisDeVries & van Swaay, 2006), while also leading to increases in plant nitrogen content and changes in host nutritional quality for insect herbivores (Vogels et al., 2023).

Thus, the impacts of two major anthropogenic stressors on insect populations largely depend on the interplay of changing host nutritional quality, vegetation structure and microclimate with insect thermal and nutritional requirements. If there is a trade-off between host plants with high nutritional quality and warm microhabitats, species requiring warmer conditions may be unable to access the potential benefits of higher host nutritional quality driven by nitrogen pollution, unless higher macroclimatic temperatures enable them to use cooler microhabitats.

Here, we investigate how host plant nutritional quality (nitrogen content) and microclimate shape habitat use by a grassland butterfly, *Lycaena phlaeas* (Small Copper), and the relationship between these two aspects of habitat quality in this system. Specifically, by studying the distribution of eggs and larvae in a *L. phlaeas* population, we assess the evidence for our hypothesised trade-off between high nutritional quality host plants and high temperatures around host plants, and how differences in vegetation structure bring this about. Female butterflies generally select host plants that maximise larval survival (Gripenberg et al., 2010; Jaenike, 1978), and the availability of hosts that meet larval requirements has consistently been shown to be the main factor constraining butterfly populations in similar systems (Bourn & Thomas, 2002; Thomas et al., 2011). Thus, we expect female oviposition choice, measured by the distribution of eggs, to indicate how *L. phlaeas* populations are constrained by thermal and nutritional requirements. By replicating sampling across seasons, we also attempt to understand how these variables are related as macroclimate changes.

Specifically, we test the following hypotheses:

1. The nutritional quality of host plants is negatively correlated with the temperature under them (microclimate), presenting a potential trade-off for *L. phlaeas* oviposition.
2. In early summer, when macroclimatic temperatures are relatively high, microclimate is less important in dictating egg-laying patterns, and host plants with higher nutritional quality are preferentially selected.
3. In autumn, when macroclimatic temperatures are lower than in early summer, *L. phlaeas* egg-laying is determined more by microclimate around host plants than their nutritional quality.

## METHODS

### Study system

*Lycaena phlaeas* is a widespread lycaenid butterfly in open habitats across Eurasia and North America. It is oligophagous on *Rumex* spp., but in north-west Europe, primarily uses *Rumex acetosa* (Common Sorrel) and *Rumex acetosella* (Sheep’s Sorrel) (Thomas & Lewington, 2014). In the United Kingdom, *L. phlaeas* undergoes a winter diapause in the larval stage and usually has between one and three generations per year depending on latitude (Thomas & Lewington, 2014). While still widespread and not categorised as threatened, it declined in abundance by 39% in the UK between 1976 and 2019 (Fox et al., 2022). Like many temperate butterfly species, *L. phlaeas* requires warm microhabitats (Streitberger et al., 2014) but will also respond strongly to changes in host nutritional quality in laboratory experiments. In oligotrophic and mesotrophic environments, *L. phlaeas* larvae appear to be nitrogen-limited, and benefit from moderate increases in plant nitrogen content, before performance declines at high nitrogen levels (Kurze et al., 2018; Langdon, 2024). Thus, they have high potential to be impacted by changes in vegetation structure, which alter microclimates, but also by changes in plant nitrogen content. Predicting the likely outcome of these changes requires an understanding of the relative strength of their thermal and nutritional constraints.

### Data collection

Our study took place at Shotover Country Park (Oxfordshire, United Kingdom, 51°44′42″N, 001°11′01″W). This site is a typical habitat for *L. phlaeas* in lowland UK, a mesotrophic grassland where the butterfly uses *R. acetosa* as its sole larval host plant and has three generations per year. *Rumex acetosa* plants were searched systematically across the study area for *L. phlaeas* eggs or larvae. For each occupied plant a paired control plant was then selected. The control plant was the nearest unoccupied *R. acetosa* plant in a randomly selected cardinal direction (based on randomly generated number between 1 and 4 corresponding to North, East, South or West respectively). For both the occupied and control plant in each pair, a 50cm quadrat with 10cm grids was used to measure the surrounding percentage cover of living vegetation, bare ground and leaf litter.

Sward height around each plant in each pair was measured following Stewart et al. (2001), using a ruler to read the height below which an estimated 80% of vegetation was growing. The area of each plant was estimated as the dimensions of the tightest-fitting rectangle, i.e. the longest dimension of the plant viewed from above, and the maximum dimension perpendicular to this. The three largest leaves without *L. phlaeas* eggs or larvae were then removed from each plant for chemical analysis, and their length and breadth measured and multiplied to approximate their area, which was averaged across the three leaves as a measure of leaf size. Leaf samples were placed in glassine envelopes, transported to the laboratory the same day and frozen at -10°C until further analysis. The microclimate of each plant was measured as the temperature at the soil surface directly under the plant in sunny conditions between 12:00 and 14:30 (the hottest part of the day, when differences between plants were expected to be most apparent and most relevant for the activity and performance of thermophilous insects) using an ultrafine thermocouple (TC Direct Ultra Fine Wire PFA type K Thermocouple 0.08mm diameter, with handheld Type K Thermocouple Indicator; Tolerance ± 1.5°C; TC Direct, Uxbridge, United Kingdom). The same thermocouple was also used to measure ambient temperature (unshielded, in full sunlight) at approximately 1.8m above the ground. This fieldwork was carried out both in early summer (10-16 June 2023) and early autumn (21-28 September 2023), when thermal constraints were expected to differ due to differences in ambient (macroclimatic) temperature.

### Chemical analysis

Cost limitations meant that we were unable to analyse the nutritional quality (in terms of nitrogen content) of every plant sample. Instead, we aimed to identify reliable proxies for plant nitrogen content that we could use in our analysis. During previous work, we observed that the area of *R. acetosa* plants and the size of their leaves appeared to increase with plant nitrogen content (Langdon, 2024). We therefore performed a principal component analysis of average leaf size and plant area across all sampled plants and separated plants into 30 groups based on the first principal component. Within these, we made a further two groups based on sampling season (i.e. 60 groups in total, one for each season by principal component group combination). From each of these we randomly selected one plant for chemical analysis. This gave a total of 58 plants (as two group x season combinations lacked plants). We expected that analysing leaf samples across the full range of plant morphology (plant area and leaf size) would provide the best chance of detecting relationships between these proxy measurements and plant nitrogen content.

Selected leaf samples were dried in individual glassine envelopes at 70°C for 48 hours before being transferred to microtubes and ground to a fine powder via bead beating (MP Bio FastPrep-24 5G, MP Biomedicals, Irvine, USA). Leaf samples were beaten for two minutes with three ceramic beads of diameter 2.3mm (Zirconia/Silica, Thistle Scientific, Warwickshire, England). Approximately 10mg of each leaf sample was then analysed for carbon and nitrogen content via combustion using a Thermo Flash Carlo Erba CN analyser (EA 1112 series) (Thermo Fisher Scientific, Waltham, Massachusetts, USA) by Forest Research (Alice Holt, United Kingdom).

### Statistical analysis

To identify which of plant area, average leaf size, or a combination of the two was the most appropriate proxy for plant nitrogen content, we ran models of plant nitrogen content as a function of average leaf size and plant area (separately and together) and the first and second principal components of their PCA (alone, but not together, since this would be the equivalent to using both unmodified variables), all with a dummy ‘season’ (summer or autumn) variable to control for differences between seasons. We compared the performance of these models via their AIC, BIC and R^2^, as well as calculating the Pearson correlation coefficients between each individual proxy and nitrogen content, looking for a combination of high explanatory power (a robust proxy) and parsimony (i.e. one variable that performs almost as well as two would be preferred, for easier interpretation of later modelling steps).

*Lycaena phlaeas* habitat use was analysed across seasons using two linear models comparing the measurements of plant temperature and host quality for occupied and unoccupied plants in early summer and autumn. For each variable we thus ran a model with the following structure:

*Variable for Plant _i_ ∼ Occupancy Status of Plant _i_ * Season in which plant _i_ was sampled*.

Autumn was used as the reference level for season, so that estimates for the seasonal interaction indicate the change in the environmental variables of occupied plants from autumn to early summer.

Temperature measurements were made over several days with varying weather conditions in each field season. To account for the effect of daily variation in ambient temperature, and its influence on the microclimate temperatures measured under plants, we therefore centred and scaled our temperature measurements by date. Thus, microclimate temperature was relative to others measured on that date, and our model indicated whether *L. phlaeas* was using the warmest plants, but not how warm these actually were. Our host nutritional quality proxy was also centred and scaled to match this. We compared the results of this model to one using the scaled predicted nitrogen content of plants instead of our proxy for this, with the inverse of the scaled standard error of predicted nitrogen content as a weight.

To test the hypothesis that host plant microclimate and nutritional quality are negatively related, we modelled the relationship between temperature and host nutritional quality using a linear model. During data collection we noted that many of the plants used for *L. phlaeas* egg-laying were in areas disturbed by *Talpa europaea* (European Mole) activity (as also observed by Streitberger et al., (2014) for *L. phlaeas* and Streitberger & Fartmann (2013) for *Pyrgus malvae*). In areas with high soil nitrogen and plant productivity, this disturbance could create bare ground and warm microclimates where we would otherwise have expected them to be absent, thereby disrupting our hypothesised relationship. We therefore included an interaction between presence or absence of bare ground and host nutritional quality in this model. As for the egg-laying model, we also ran this model using the scaled predicted nitrogen content of plants instead of our proxy measure, with the inverse of the scaled standard error of predicted nitrogen content as a weight.

To understand what might drive the hypothesised relationship between temperature and host nutritional quality we then examined the effect of environmental variables (bare ground, vegetation cover, leaf litter cover and sward height) on relative microclimate temperature using linear models. Finally, we tested the effect of sward height and litter cover on host nutritional quality in the same way (omitting bare ground and vegetation cover and dropping plants with bare ground from the analysis since these mostly reflected *T. europaea* disturbance rather than underlying soil conditions).

All analyses were carried out in R (R Core Team, 2024) using the ‘tidyverse’, ‘readr’ and ‘Hmisc’ packages for data manipulation (Harrel, 2024; Wickham et al., 2019, 2024), ‘performance’ for model comparison (Lüdecke et al., 2021), and ‘ggplot2’, ‘cowplot’ and ‘ggeffects’ for plotting (Lüdecke, 2018; Wickham, 2016; Wilke, 2024). Model fit was assessed via inspection of plots of the residuals and fitted values.

## RESULTS

### Host nutritional quality and proxies

Among our 58 *R. acetosa* leaf samples, nitrogen content (by mass) averaged 2.54% ±0.10 with a maximum of 4.39% and a minimum of 0.98%. This translated into molar content averaging 1815.79 micromoles g^-1^ ±70.78 with a maximum of 3130.64 micromoles g^-1^ and a minimum of 701.09 micromoles g^-1^. This led to molar C:N ratios of between 11.24 and 51.98, averaging 22.41 ±1.01, which is higher than the optimal value for *L. phlaeas* larvae of 18.1 estimated in previous research (Langdon, 2024). The distribution of these measurements is plotted in Figure S1, although the stratified nature of our sampling means that these are not representative of the whole population of *R. acetosa* at the study site.

Plant area and leaf size were positively correlated (Pearson’s R = 0.61, t = 13.54, df = 316, p < 0.001) and the first principal component of their PCA explained 79.25% of the variance in the two measurements (Table 1). Models of plant nitrogen content as a function of these proxies and principal components (results in Table S1 and Figure S2) indicated that leaf size was the best proxy for nitrogen content. Of the individual options, it showed the strongest positive correlation with plant nitrogen content (Table 2), and models with just leaf size performed better than either principal component by themselves (Table 3). The model with leaf size and plant area performed only marginally better than leaf size alone (Table 3). Therefore, on the basis of parsimony and predictive power, leaf size was chosen as our proxy for plant nitrogen content. The results were qualitatively the same when C:N ratio was used instead of N content as the ‘true’ measure of host nutritional quality (Table S2 and Table S3).

**Table 1:** Results of Principal Components analysis of host nutritional quality proxies.

|  | **PC1** | **PC2** |
| --- | --- | --- |
| Eigenvalue | 1.59 | 0.42 |
| % Variance Explained | 79.25 | 20.75 |
| Average Leaf Size (mm<sup>2</sup>) loading | -0.71 | -0.71 |
| Plant Area (cm<sup>2</sup>) loading | -0.71 | 0.71 |

**Table 2:**
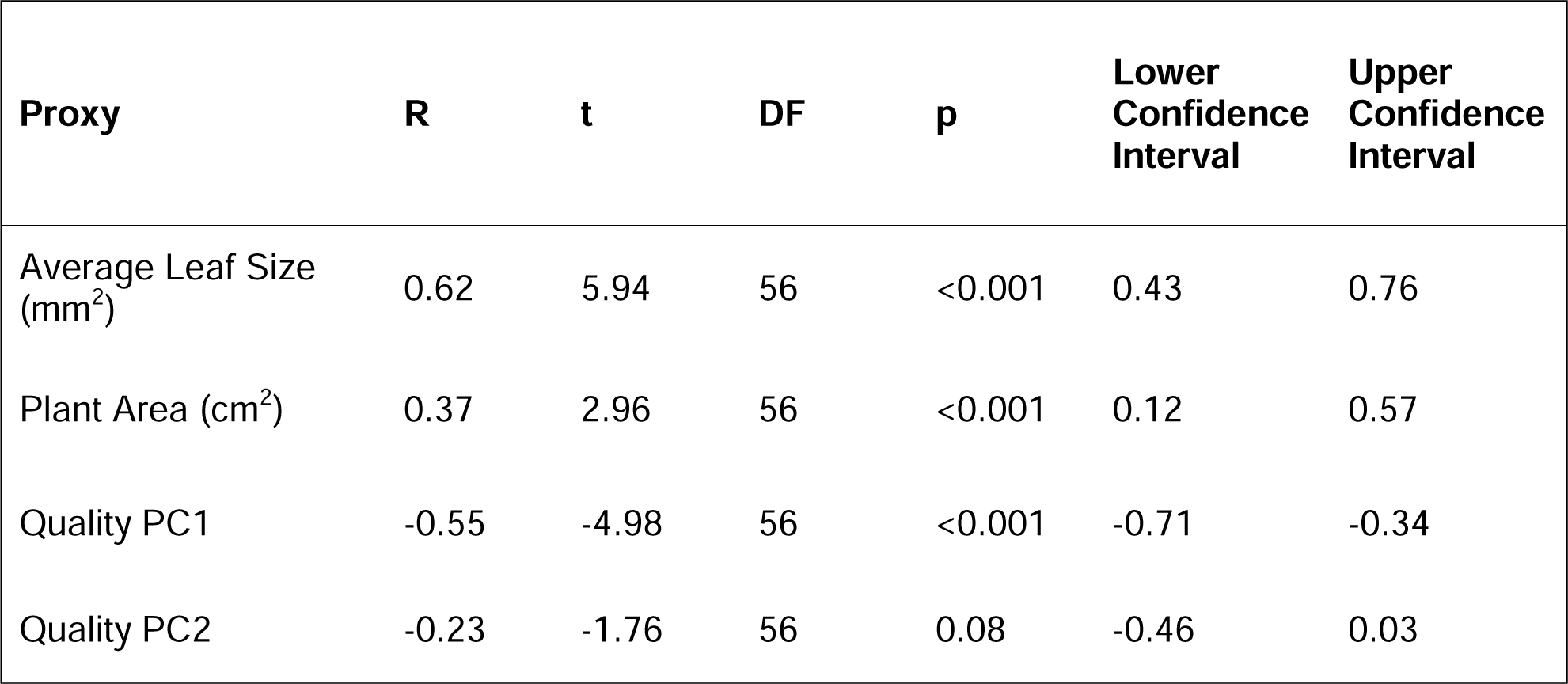
Pearson’s correlation coefficients and significance tests for correlation of possible plant proxies with plant nitrogen content (measured in micromoles g^-1^).

| Proxy | R | t | DF | p | Lower<br>Confidence<br>Interval | Upper<br>Confidence<br>Interval |
| --- | --- | --- | --- | --- | --- | --- |
| Average Leaf Size (mm<sup>2</sup>) | 0.62 | 5.94 | 56 | <0.001 | 0.43 | 0.76 |
| Plant Area (cm<sup>2</sup>) | 0.37 | 2.96 | 56 | <0.001 | 0.12 | 0.57 |
| Quality PC1 | -0.55 | -4.98 | 56 | <0.001 | -0.71 | -0.34 |
| Quality PC2 | -0.23 | -1.76 | 56 | 0.08 | -0.46 | 0.03 |

**Table 3:** Comparison of models relating plant nitrogen content to different possible proxies.

| Proxies | AIC | BIC | R<sup>2</sup> |
| --- | --- | --- | --- |
| Average Leaf Size (mm<sup>2</sup>) + Plant Area (cm<sup>2</sup>) | 873 | 883 | 0.41 |
| Average Leaf Size (mm<sup>2</sup>) | 871 | 879 | 0.41 |
| Plant Area (cm<sup>2</sup>) | 883 | 891 | 0.27 |
| Quality PC1 | 874 | 882 | 0.37 |
| Quality PC2 | 892 | 900 | 0.15 |

### Microclimate under *R. acetosa* plants

Ambient temperatures in early summer averaged 24.7°C ±0.1, and in autumn 16.3°C ±0.1, approximately 8.4°C lower (t = 60.42, df = 185.49, p < 0.001) (Figure S3). Matching this, microclimate temperatures measured under *R. acetosa* plants were higher in early summer, averaging 36.7°C ±0.5 compared to 18.4°C ±0.1 in autumn, a difference of 18.3°C that was also statistically significant (t = 38.34, df = 217.53, p < 0.001) (Figure S3). The maximum temperature recorded under *R. acetosa* plants in early summer was 56°C and in autumn 23°C.

### Impacts of host plant nutritional quality and microclimate on *L. phlaeas* egg-laying

Overall, 318 plants were sampled, 184 in early summer and 134 in the autumn (evenly divided into occupied plants and control plants). Occupied host plants were significantly warmer than unoccupied host plants (β = 1.22 ± 0.14, t = 8.83, df = 314, p < 0.001), but there was no difference in their nutritional quality (β = 0.15 ± 0.16, t = 0.89, df = 314, p = 0.37) (Figure 1, Table 4). This pattern was observed in both early summer and autumn, and the differences in thermal and nutritional quality of occupied and unoccupied plants were constant across seasons (Figure 1, Table 4). Results for plant quality were qualitatively the same when scaled predicted nitrogen content of plants was used instead of the proxy of leaf size (Table S4).

**Figure 1:**
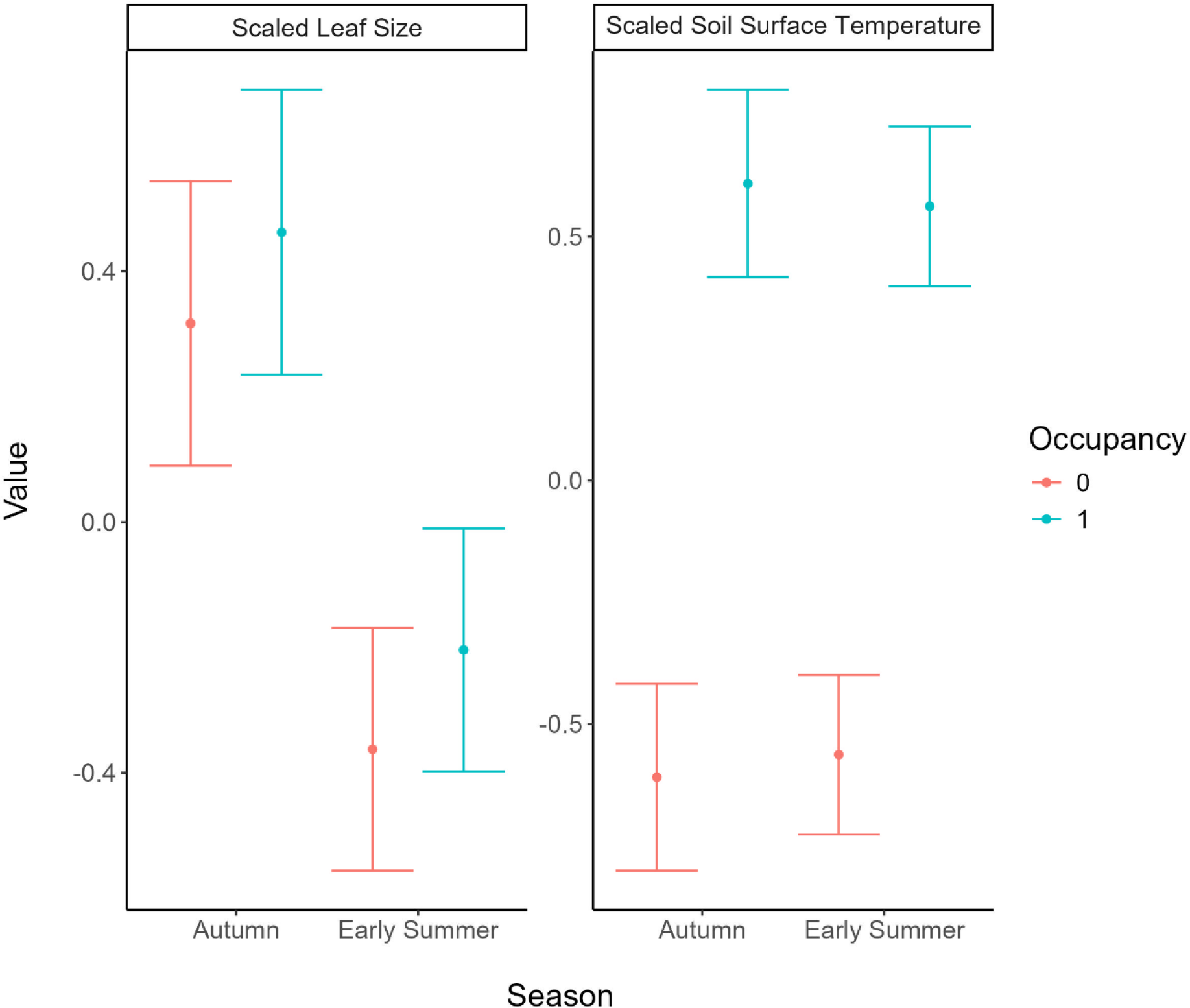
Mean values of scaled leaf size and soil surface temperature measured on and around occupied and unoccupied host plants of *L. phlaeas* in early summer and autumn. Points show predicted marginal effects from linear models (Table 4) and bars their 95% confidence intervals.

**Table 4:** Outputs from models assessing differences in nutritional quality (scaled leaf size) and microclimate (scaled soil surface temperature) of occupied and unoccupied *L. phlaeas* host plants in early summer and autumn.

| Measurement | Term | Estimate | Standard Error | t | p |
| --- | --- | --- | --- | --- | --- |
| Scaled Leaf Size | Intercept | 0.32 | 0.12 |  |  |
|  | Occupied | 0.15 | 0.16 | 0.89 | 0.37 |
|  | Season | -0.68 | 0.15 | -4.48 | <0.001 |
|  | Occupied:Season | 0.01 | 0.21 | 0.06 | 0.95 |
| Scaled soil surface temperature | Intercept | -0.61 | 0.1 |  |  |
|  | Occupied | 1.22 | 0.14 | 8.83 | <0.001 |
|  | Season | 0.05 | 0.13 | 0.36 | 0.72 |
|  | Occupied:Season | -0.09 | 0.18 | -0.51 | 0.61 |

On occupied plants, the mean number of eggs in early summer was 2.32 ±0.29 with a maximum of 16 and a total (across all plants) of 213. In autumn, these figures were 1.90 ±0.29, 8 and 127 eggs respectively. The median in both seasons was 1 egg.

### Relationships between microclimate and host nutritional quality

As expected, in the absence of bare ground, host nutritional quality (scaled leaf size) was significantly negatively related to scaled soil surface temperature under plants (β = -0.30 ± 0.08, t = -3.94, df = 313, p < 0.001). At the lowest leaf sizes, plants with bare ground were significantly warmer than those without bare ground (β = 0.80 ± 0.11, t = 7.13, df = 313, p < 0.001), and the presence of bare ground (associated with *T. europaea* activity) made the slope of the relationship between host nutritional quality and scaled temperature under plants significantly less negative, almost entirely flattening it (β = 0.28 ± 0.11, t = 2.62, df = 313, p < 0.01) (Figure 2). Results did not qualitatively differ when using predicted nitrogen content as our host nutritional quality variable (see Table S5).

**Figure 2:**
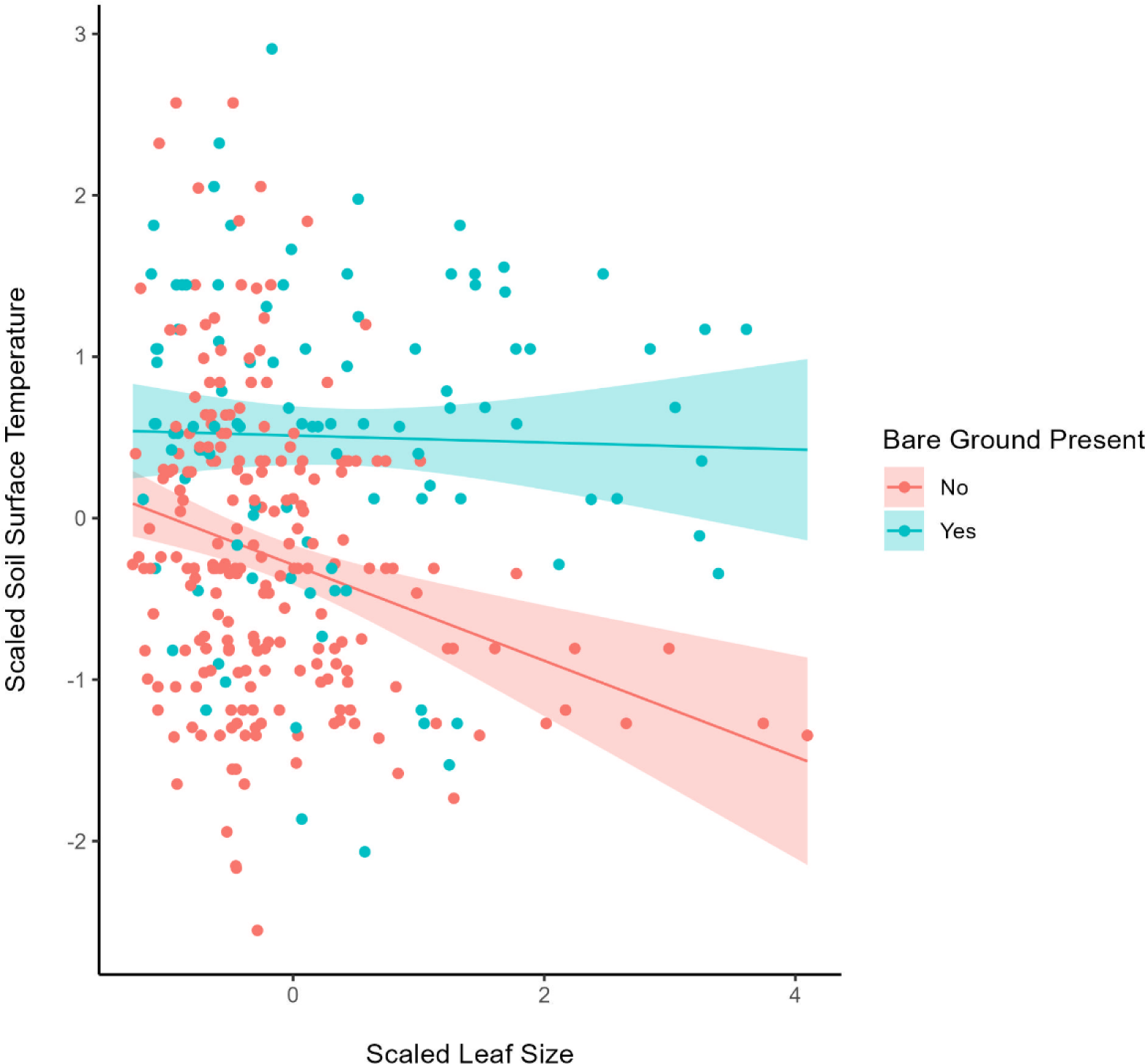
The modelled relationship between host nutritional quality (measured using the proxy of leaf size), microclimate and the presence or absence of bare ground. Predictions from linear model plotted with their 95% confidence intervals and the underlying data.

### Relationships between microclimate and vegetation structure

Variables characterising vegetation structure around *R. acetosa* plants were all significantly associated with scaled soil surface temperature in the manner expected (Figure 3). Cover of bare ground (β = 0.020 ± 0.0028, t = 7.15, df = 315, p < 0.001) and cover of litter (β = 0.018 ± 0.0046, t = 3.95, df = 315, p < 0.001) were positively associated with scaled soil surface temperature, and sward height (β = -0.037 ± 0.0036, t = -10.33, df = 314, p < 0.001) and cover of vegetation (β = -0.024 ± 0.0037, t = -6.50, df = 315, p < 0.001) were negatively associated. Individually, sward height explained more variation in microclimate than any other variable (R^2^ = 0.25), followed by bare ground (R^2^ = 0.14), vegetation cover (R^2^ = 0.12) and litter cover (R^2^ = 0.044).

**Figure 3:**
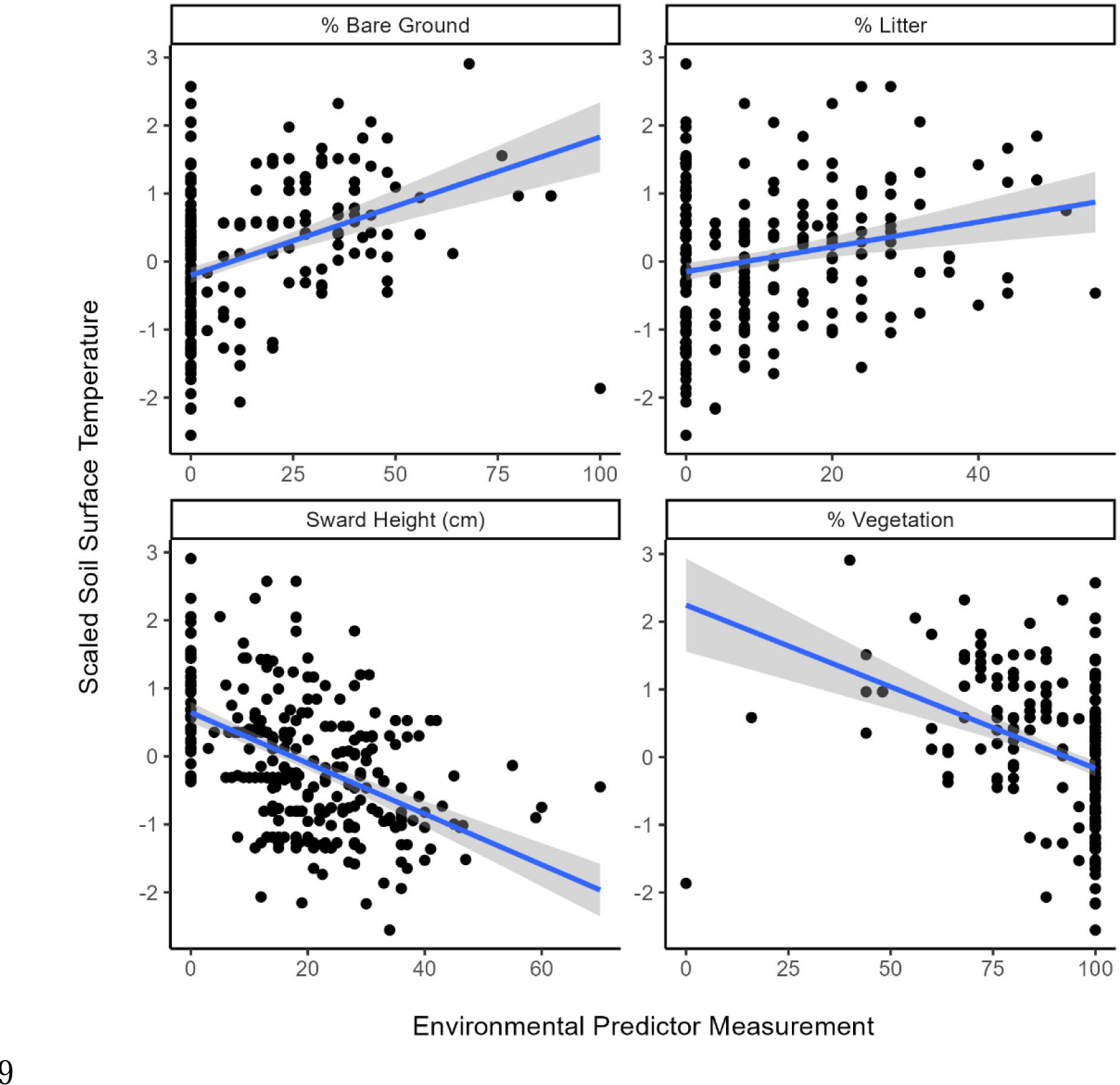
Relationships between vegetation structure around host plants and the relative temperatures measured under them. Predicted trends from linear models are shown with their 95% confidence intervals and the underlying data.

### Relationships between vegetation structure and host nutritional quality

Litter cover and sward height were both related to our host nutritional quality proxy (average leaf size) in the manner expected, the former significantly negatively associated (β = -9.14 ± 2.00, t = -4.56, df = 212, p < 0.001) and the latter positively (but not significantly) associated (β = 0.272 ± 2.52, t = 0.11, df = 211, p = 0.91). While both litter cover and sward height were poor predictors of host nutritional quality, the litter model explained more variation in host nutritional quality than sward height (litter R^2^ = 0.085, sward height R^2^ = 0.0000055).

## DISCUSSION

Insect herbivores are expected to be constrained by both the nutritional quality and the microclimate of their host plants. We predicted (hypothesis 1) that, in open habitats, insects may face a trade-off between these constraints, due to plants with high nutritional quality (nitrogen content) being expected to occur in cooler microclimates. In our study of egg-laying patterns of the widespread butterfly *L. phlaeas*, we found strong support for this hypothesis of a significant negative relationship between host plant nitrogen content and microclimate (although this relationship was fundamentally altered if *T. europaea* activity generated bare ground close to host plants). However, we found no evidence that the butterfly’s egg-laying decisions shift between seasons to favour host plant nutritional quality when thermal constraints are eased by higher ambient temperatures (our hypotheses 2 and 3). Egg-laying of *L. phlaeas* on *R. acetosa* was primarily determined by microclimate. We found no significant effect of host nutritional quality (proxies of nitrogen content) on the probability of egg-laying, and no difference between a warmer period (in early summer) and a cooler one (in early autumn).

Given that most plants in our sample appeared to be of sub-optimal nutritional quality for *L. phlaeas* (Figure S4) (Langdon, 2024), and evidence that other butterfly species are able to select host plants on the basis of nutritional quality (Bourn & Thomas, 1993; Letourneau & Fox, 1989; Prudic et al., 2005), it appears that female *L. phlaeas* are missing an opportunity to maximise larval fitness by laying their eggs on the minority of plants with higher nutritional quality. Instead, microclimate appears to be the dominant driver of oviposition choice and, unless there happened to be soil disturbance under a host plant, plants with warmer microclimates were of lower nutritional quality (as hypothesised).

It therefore appears that in our system ovipositing *L. phlaeas* face a choice between high nutritional quality host plants or warm host plants, adding to previous work showing that egg-laying butterflies may have to balance nutritional requirements against thermal requirements (Krämer et al., 2012). The apparent prioritisation of the latter implies that thermal constraints are strong for *L. phlaeas*, in line with previous work (León-Cortés et al., 2000; Streitberger et al., 2014) and override nutritional constraints. Many other studies have highlighted the importance of thermal constraints on habitat use by butterfly species but without examining host plant nutritional quality (e.g. Curtis & Isaac, 2015; Johansson et al., 2024; Roy & Thomas, 2003; Thomas et al., 1986; C.D. Thomas et al., 2001; Wilby et al., 2024). Although we do not have data relating *L. phlaeas* larval performance to temperature, previous experimental work (Langdon, 2024) suggests that although most *R. acetosa* plants in our system were of sub-optimal nutritional quality for *L. phlaeas* (Figure S4), they were still in a range where larval survival was around 80% of the maximum. Furthermore, by feeding selectively on different plants or plant parts to regulate their nutrient intake, larvae in our study system might be able to buffer the impacts of low host nutritional quality. Thus, the fitness consequences of using hosts with low nutritional quality at our study site are likely to be relatively small for *L. phlaeas*. A further intriguing possibility is that the fitness costs associated with sub-optimal food quality may also be reduced by high temperatures. Clissold et al. (2013) found that the efficiency of macronutrient absorption varied with temperature for *Locusta migratoria* nymphs, and that protein absorption was more efficient at higher temperatures for nymphs feeding on *Themeda triandra*. Protein is the main pool of nitrogen in plants, suggesting that increases in absorption efficiency on warm host plants could offset the cost of their lower nitrogen levels. Similar work is lacking for Lepdioptera, however, and Clissold et al. (2013) did not observe the same trend in locusts fed *Triticum aestivium*, suggesting that such increases in absorption efficiency are unlikely to be universal.

In spite of the far higher ambient temperatures and microclimatic temperatures around *R. acetosa* in early summer, we found no support for our hypothesis that *L. phlaeas* would shift oviposition choices to use hosts with higher nutritional quality in relatively cooler microclimates in early summer compared to autumn. This contrasts with previous work on another lycaenid butterfly, *Polyommatus bellargus*, in which higher ambient temperatures in early summer allowed the use of larval host plants growing in cooler microhabitats, doubling habitat availability for the species (Roy & Thomas, 2003). This difference is difficult to explain and seems unlikely to reflect a difference in thermal constraints given that *L. phlaeas* occurs north into the Arctic Circle (Kudrna et al., 2011) while *P. bellargus* reaches the northern edge of its range in southern England, where it is dependent on conservation management to provide suitably warm microhabitats (Thomas, 1983).

At our study site, the cooling effect of the tall swards (mean sward height = 17.14 cm, ±0.75) may have meant that only a small proportion of *R. acetosa* plants were thermally suitable for *L. phlaeas*. In such mesotrophic grasslands, *L. phlaeas* may breed mainly in small habitat islands of disturbed ground or low fertility soil where host plant microclimates are suitably warm. Even when ambient temperatures are high, e.g. in early summer, and thermal constraints are expected to be lessened, there may be little other suitable habitat for the butterfly to use because most host plants are still too cool. Indeed, we often observed that egg-laying took place on plants growing on bare ground generated by *T. europaea* disturbance, which was positively associated with the temperature under host plants. In fact, 55% of plants with *L. phlaeas* eggs had some bare ground within 25cm of them, compared to only 11% of unoccupied plants. Streitberger et al. (2014) also observed high rates of host plant use by *L. phlaeas* on *T. europaea* earthworks, while Streitberger & Fartmann (2013) found the same for the hesperiid *Pyrgus malvae*. Another skipper butterfly, *Hesperia comma*, has also been shown to favour *Lasius flavus* (Yellow Meadow Ant) mounds for oviposition (Streitberger & Fartmann, 2016), while soil disturbance caused by foraging behaviour of *Sus scrofa* (Wild Boar) has similarly been positively linked to butterfly oviposition choice by *P. malvae* (de Schaetzen et al. 2018) and two fritillary species (Scherer et al., 2025). Our results thus add to the growing body of evidence demonstrating the importance of small-scale soil disturbance for creating microhabitat heterogeneity that contributes to the maintenance of diverse grassland plant and insect communities (e.g. Seifan et al., 2010; Fleischer et al., 2013).

The dependence of *L. phlaeas* on small islands of warm microhabitat may explain why the response to seasonal variation in temperature that we documented differs from that found for *P. bellargus* (Roy & Thomas, 2003), where habitats often have uniformly high thermal suitability as a result of heavy grazing. It also reinforces the importance of continued conservation management of open habitats to create suitable microclimates for thermally-constrained species, in spite of the potential benefits of climate change to these species (O’Connor et al., 2014). In a broader sense, habitat management to create or maintain microclimatic heterogeneity will help to buffer a wide variety of species from the impacts of extreme weather events (Hayes et al., 2024) and climate change (Suggitt et al., 2018).

We expected that higher nitrogen content in host plants would reflect higher soil nitrogen levels, which would in turn enhance plant productivity, leading to denser vegetation and lower temperatures around host plants. We found some support for this mechanism, with strong associations between measures of vegetation structure and temperature (shorter, more open vegetation meant higher temperatures) but only weak relationships in the directions expected between vegetation structure and host nutritional quality – higher litter cover was significantly, albeit weakly, associated with lower nutritional quality, while higher sward height was positively, but not significantly, related to host nutritional quality. Despite this, it is difficult to envisage another mechanism that could alter host nutritional quality and vegetation structure to affect microclimate simultaneously. Grazing or cutting often does the latter, but the most commonly-reported plant response to grazing appears to be an increase in nitrogen uptake and availability in young plant tissues (e.g. Jaramillo & Detling, 1988; Ruess, 1984), which would lead to a positive correlation between temperature and host nutritional quality, rather than the inverse as observed here. The rather weak relationships that we detected between host nutritional quality and vegetation structure may reflect the fact that our measures of vegetation structure, though commonly used approximations, imperfectly capture the factors driving the link between host nutritional quality and microclimate. Further work relating a more detailed characterisation of vegetation structure to host nutritional quality and, crucially, measuring underlying soil nitrogen levels would help to establish whether this trade-off is indeed a general phenomenon for grassland insects.

Our results indicate that habitat use by *L. phlaeas* is driven by thermal requirements and not by nutritional requirements, and egg-laying females may be forced to use host plants of suboptimal nutritional quality. This provides useful context for laboratory experiments showing strong impacts of plant nitrogen content on the performance of *L. phlaeas*, with negative implications for fitness and survival under levels of nitrogen pollution associated with management of agricultural grasslands (Kurze et al., 2018). Our results suggest that such responses may not scale up to impacts of nitrogen pollution on *L. phlaeas* populations in the field via changes in plant nutritional quality, because high-nitrogen plants will be thermally unsuitable. Thus, *L. phlaeas* and species with similar thermal requirements seem more likely to be negatively impacted by changes to vegetation structure under nitrogen pollution, than via changes in host nutritional quality (Nijssen et al., 2017).

It is important to note that our study took place in the United Kingdom, near the edge of the species’ European range. Thus, our results are not necessarily generalisable across the full range of *L. phlaeas*; in warmer areas, the species may no longer be constrained to prioritise microclimate over nutritional quality. However, we did not observe a shift in egg-laying from warmer host plants in autumn to cooler, higher quality host plants in early summer, despite ambient midday temperatures being 8.4°C higher on average. This suggests that climatic differences would need to be substantial to release the butterfly from the constraint.

Our results align with previous work by Klop et al. (2015), who performed a similar study with the grassland butterfly *Lasiommata megera,* assessing larval microhabitat requirements in the field and responses to plant nitrogen content under laboratory conditions. They found only positive effects of fertilisation, and a tendency to use warmer microhabitats, suggesting that the negative effects of atmospheric nitrogen deposition on *L. megera* population trends were likely to reflect losses of suitable microhabitats, offsetting any improvement in host quality. Similarly, Yang et al. (2017) tested the response of larvae of the moth *Gynaephora alpherakjj* to nitrogen fertilisation of foodplants in alpine grassland and found that growth rate and feeding rate declined with fertilisation in the field, despite positive impacts in the laboratory, attributing this difference to the lower temperatures at ground level in fertilised plots. Interestingly, however, we found that soil disturbance by *T. europaea* disrupted the trade-off between host nutritional quality and temperature around plants (by creating high temperatures around plants of high nutritional quality), suggesting that soil disturbance generated in this manner, or at larger scales by grazing animals, may contribute to maintaining insect herbivore species of early-successional habitats under nitrogen pollution.

In conclusion, we found that *L. phlaeas* was strongly thermally constrained, and selection of larval host plants for egg-laying was determined by warm microclimates rather than host nutritional quality, even during warmer periods of the year. Indeed, these constraints may force *L. phlaeas* to use plants of lower nutritional quality, as we observed a negative association between host nutritional quality and temperatures around plants. We suggest this could be a general trade-off faced by grassland insect species, driven by underlying variation in soil nitrogen and its impact on host nutritional quality and vegetation structure, but further investigation is needed to confirm the mechanisms underlying this interesting pattern. Overall, this places the strong responses of herbivores, such as *L. phlaeas*, to variation in host nutritional quality in fertilization experiments into context; under field conditions, thermophilous insects are more likely to be negatively impacted by nitrogen pollution via changes in vegetation structure than positively via increased host nutritional quality. Our results suggest that habitat management to maintain or create warm microhabitats will be essential to the conservation of this declining butterfly, and other thermophilous insects, even under a changing climate.

## Acknowledgements

We would like to thank Julian Cooper (Oxford District Council) for permission to carry out fieldwork at Shotover Country Park and François Bouchereau (Forest Research) for plant chemistry analyses. Lisbeth Hordley generously provided statistical advice.

## Author contributions

**Willam B.V. Langdon** Conceptualization (equal); Investigation (lead); Formal Analysis (lead); Writing – Original Draft Preparation (lead); Writing – Review & Editing (equal). **Richard Fox** Conceptualization (supporting); Supervision (equal); Writing – Original Draft Preparation (supporting); Writing – Review & Editing (equal). **Owen T. Lewis** Conceptualization (equal); Supervision (equal); Writing – Review & Editing (equal).

## Data availability statement

Data used in this manuscript are available from the Dryad Digital Repository.

Langdon, W.B.V., Fox, R., Lewis, O.T. 2024. Dryad Digital Repository DOI:10.5061/dryad.hqbzkh1wk

