## Supporting Information for "Host plant use is driven by microclimate not nutritional quality in a grassland butterfly"

**Host nutritional quality**

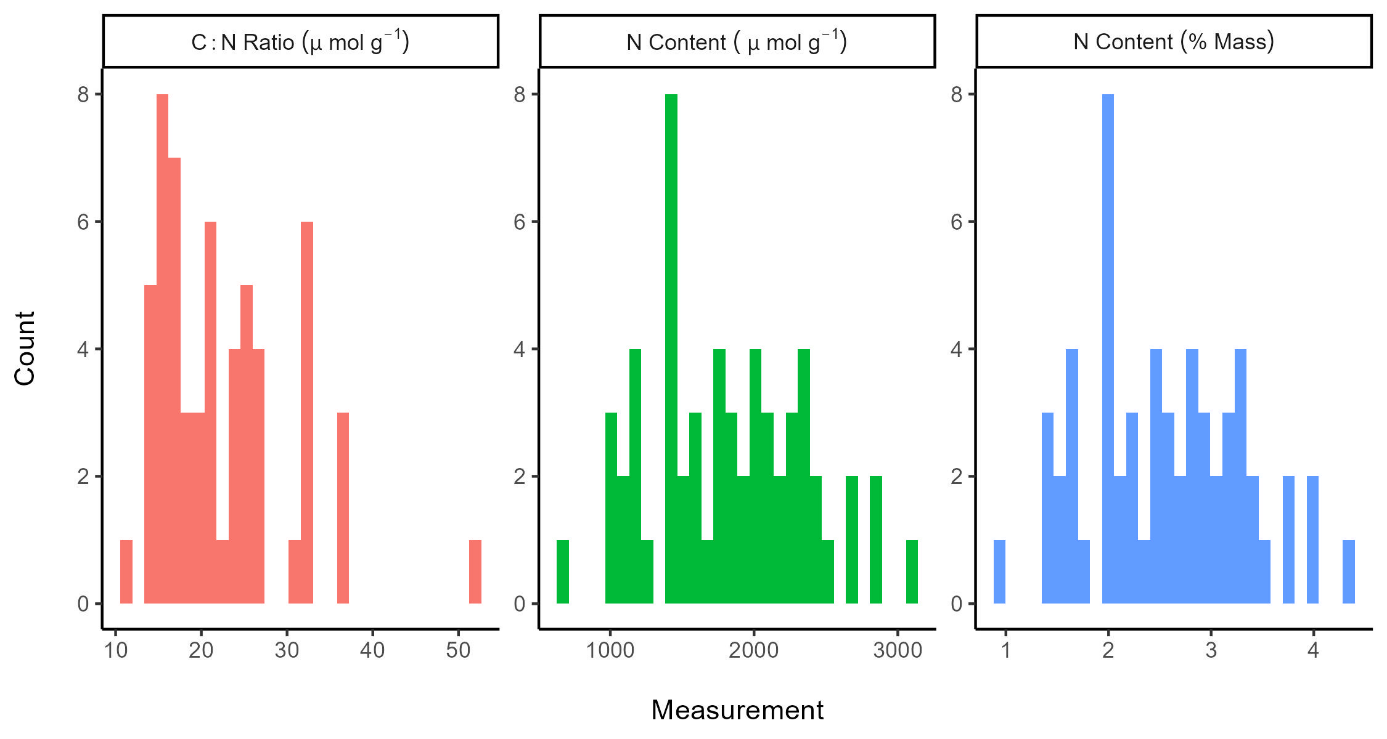

**Figure S1:** Distribution of chemical measurements for *R. acetosa* plants that were analysed. As these were a stratified sample they do not represent the true distribution in the population.

**Model results relating plant nitrogen content to nutritional quality proxies**

**Table S1:** Results of linear models relating plant nitrogen content (micromoles g^-1^) to the possible proxies assessed.

| **Model** | **Term** | **β** | **Standard Error** | **t** | **p** |
| --- | --- | --- | --- | --- | --- |
| Leaf Size + Plant Area | Intercept | 1392.44 | 142.58 | 9.77 | <0.001 |
|  | Average Leaf Size (mm^2^) | 0.42 | 0.12 | 3.58 | <0.001 |
|  | Plant Area (cm^2^) | 0.21 | 0.37 | 0.57 | 0.57 |
|  | Season | -185.18 | 127.36 | -1.45 | 0.15 |
| Leaf Size | Intercept | 1395.56 | 141.6 | 9.86 | <0.001 |
|  | Average Leaf Size (mm^2^) | 0.46 | 0.09 | 4.93 | <0.001 |
|  | Season | -166.61 | 122.42 | -1.36 | 0.18 |
| Plant Area | Intercept | 1729.49 | 118 | 14.66 | <0.001 |
|  | Plant Area (cm^2^) | 1.01 | 0.33 | 3.08 | <0.01 |
|  | Season | -396.51 | 124.38 | -3.19 | <0.01 |
| Quality PC1 | Intercept | 1754.16 | 92.59 | 18.95 | <0.001 |
|  | Quality PC1 | -147.52 | 33.03 | -4.47 | <0.001 |
|  | Season | -283.05 | 118.74 | -2.38 | <0.05 |
| Quality PC2 | Intercept | 1980.99 | 89.16 | 22.22 | <0.001 |
|  | Quality PC2 | -57.47 | 75.69 | -0.76 | 0.45 |
|  | Season | -368.19 | 144.44 | -2.55 | <0.05 |

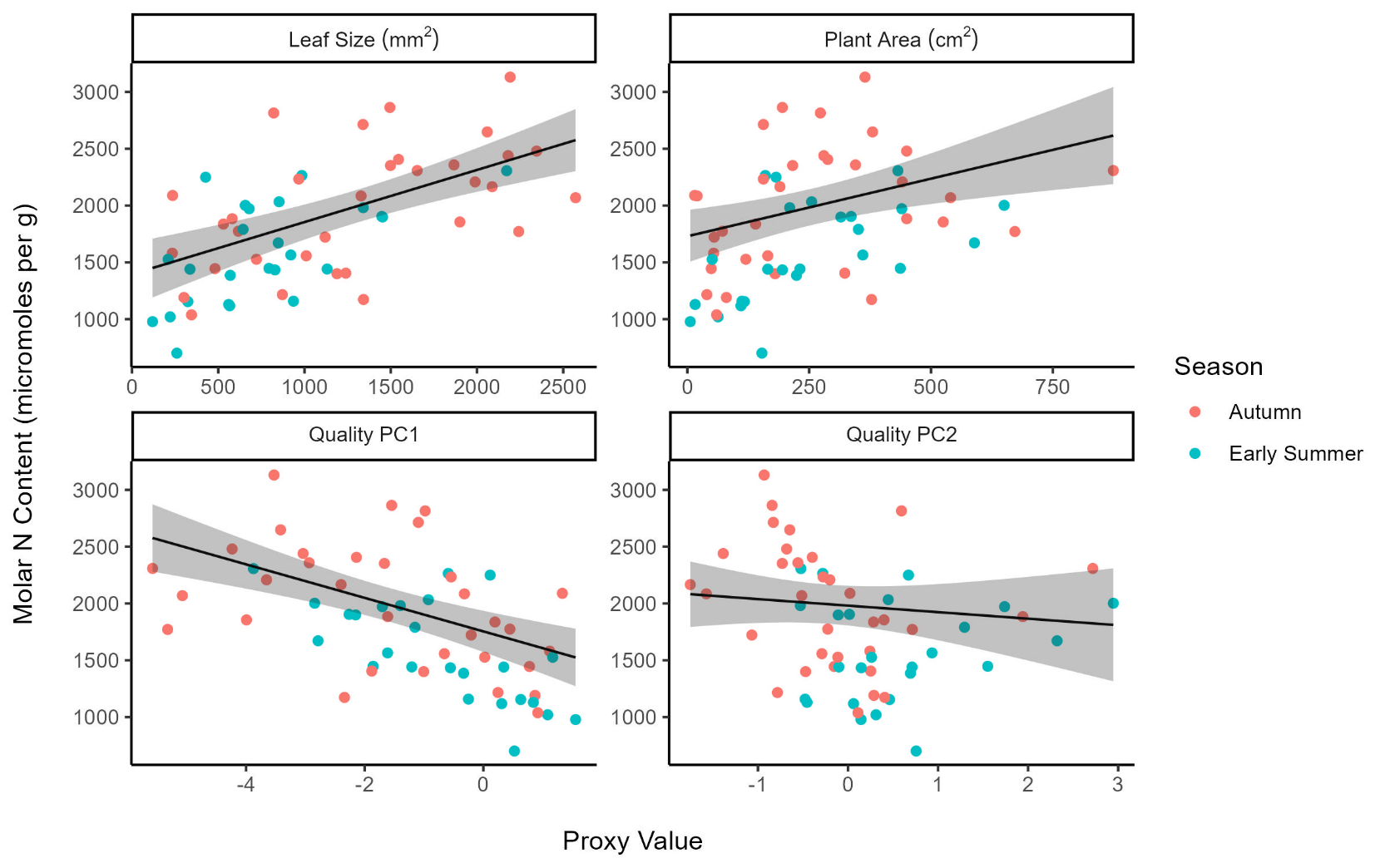

**Figure S2:** Predicted relationships between host nutritional quality proxies and plant nitrogen content. Predictions are for autumn and plotted with 95% confidence intervals from models (results Table S1) alongside the underlying data.

**Model results relating plant C:N ratio to nutritional quality proxies**

These were qualitatively the same as for plant N content (micromoles g^-1^) (Table 1 and Table 2), with average leaf size the best proxy, but plant size adding a small amount of additional explanatory power.

**Table S2:** Pearson’s correlation coefficients and significance tests for correlation of possible plant proxies with *R. acetosa* C:N ratio (measured in micromoles g^-1^).

| **Proxy** | **R** | **t** | **DF** | **p** | **Lower Confidence Interval** | **Upper Confidence Interval** |
| --- | --- | --- | --- | --- | --- | --- |
| Average Leaf Size (mm^2^) | -0.61 | -5.83 | 56 | <0.001 | -0.75 | -0.42 |
| Plant Area (cm^2^) | -0.41 | -3.38 | 56 | <0.001 | -0.61 | -0.17 |
| Quality PC1 | 0.58 | 5.27 | 56 | <0.001 | 0.37 | 0.73 |
| Quality PC2 | 0.17 | 1.32 | 56 | 0.19 | -0.09 | 0.41 |

**Table S3:** Comparison of models relating plant C:N ratio to different possible proxies.

| **Proxies** | **AIC** | **BIC** | **R^2^** |
| --- | --- | --- | --- |
| Average Leaf Size (mm^2^) and Plant Area (cm^2^) | 380 | 391 | 0.4 |
| Average Leaf Size (mm^2^) | 379 | 388 | 0.388 |
| Plant Area (cm^2^) | 389 | 397 | 0.276 |
| Quality PC1 | 381 | 389 | 0.376 |
| Quality PC2 | 401 | 409 | 0.118 |

**Microclimate under *R. acetosa* plants**

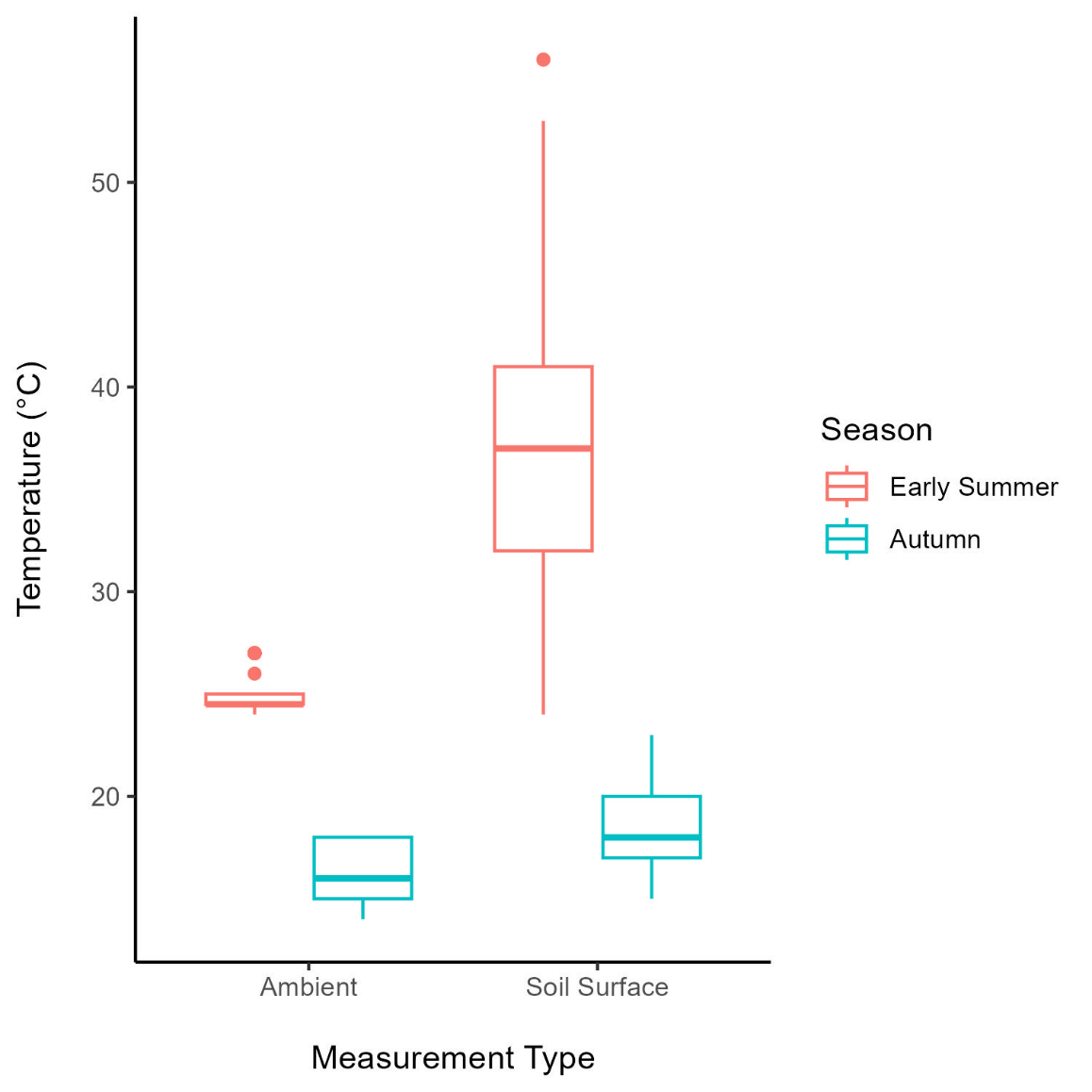

**Figure S3:** Differences in ambient temperatures and microclimate temperatures under *R. acetosa* plants measured in early summer and autumn. Boxes show the first, second (median) and third quartiles for each measurement, whiskers 1.5 x IQR above the third quartile, and below the first quartile.

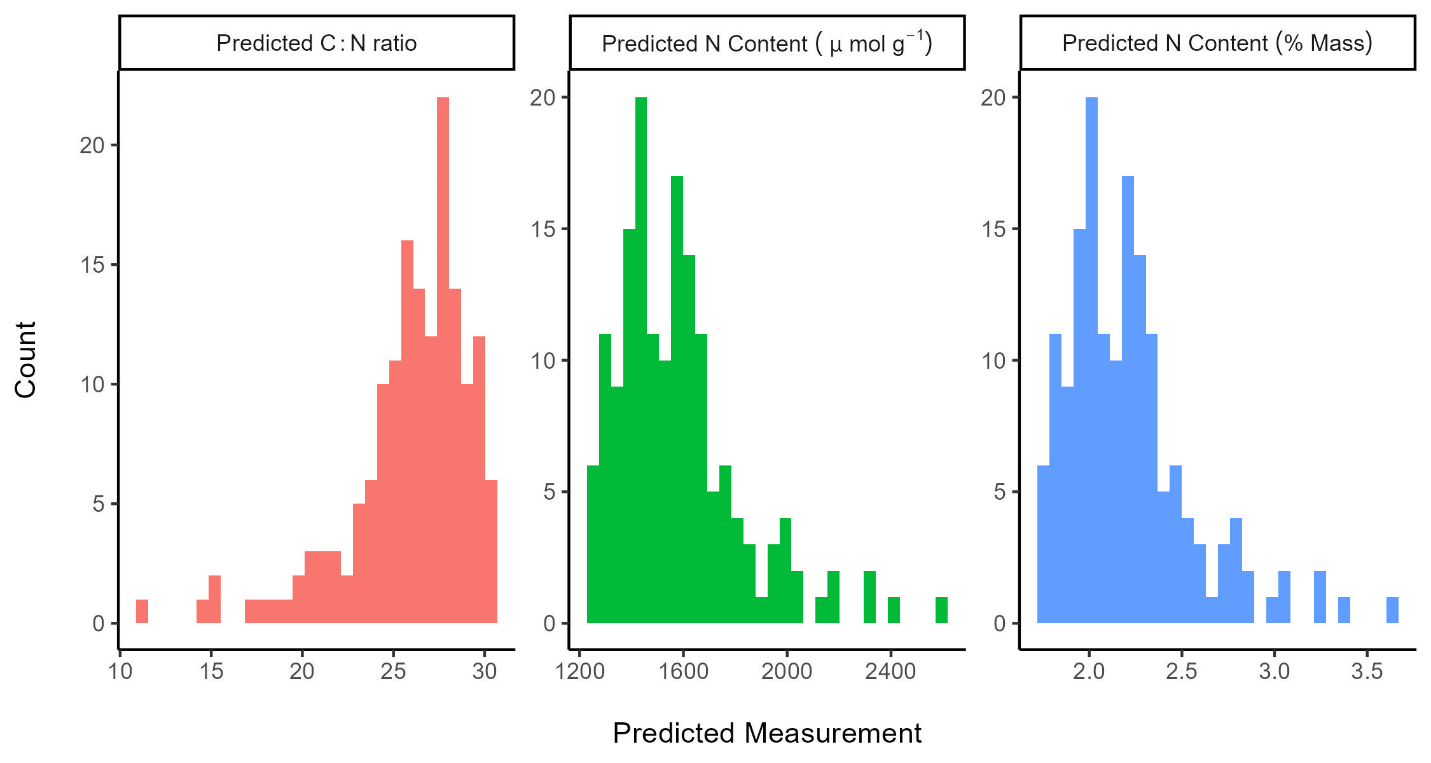
**Predicted quality of *Rumex acetosa* plants**

**Figure S4:** Predicted C:N ratio and nitrogen contents of *R. acetosa* plants at Shotover Country Park, based on our models of host nutritional quality as a function of average leaf size and plant area. These are plotted for control plants and therefore give a better sense of the likely distribution of measurements within the population, unlike figure S1 which shows a sample of plants stratified by proxies of nitrogen content, therefore giving a far more uniform distribution than is actually present. This shows that most plants were likely to be of low nutritional quality, with larvae suffering from nitrogen deficiency, as most plants are predicted to have C:N ratios higher than the value of 18.1 predicted to be optimal for *L. phlaeas* by Langdon (2024).

**Models using predicted plant nitrogen content as a measure of host quality, rather than average leaf size**

**Egg-laying model**

**Table S4:** Outputs from model assessing differences in scaled predicted nitrogen content of occupied and unoccupied *L. phlaeas* host plants in early summer and autumn. To account for uncertainty around predictions, 1/scaled standard error around predicted nitrogen content was used to weight data points in this model.

| **Term** | **Estimate** | **Standard Error** | **t** | **p** |
| --- | --- | --- | --- | --- |
| Intercept | 0.72 | 0.09 |  |  |
| Occupied | 0.13 | 0.13 | 1.01 | 0.31 |
| Season | -1.32 | 0.12 | -11.21 | <0.001 |
| Occupied:Season | -0.03 | 0.17 | -0.17 | 0.86 |

**Trade-off Model**

**Table S5:** Results of model of scaled temperature under *R. acetosa* plants as a function of their scaled predicted nitrogen content and the presence of bare ground (and 1/scaled standard error around predicted nitrogen content as weights). These were qualitatively the same as when we used our proxy for nitrogen content (leaf size) (Figure 2).

| **Term** | **Estimate** | **Standard Error** | **t** | **p** |
| --- | --- | --- | --- | --- |
| Intercept | -0.29 | 0.06 |  |  |
| Scaled N content | -0.29 | 0.07 | -3.99 | <0.001 |
| Bare Ground Present | 0.76 | 0.11 | 6.67 | <0.001 |
| Scaled N content : Bare Ground Present | 0.33 | 0.11 | 3.09 | <0.001 |
